# TFIIS is required for reproductive development and thermal adaptation in barley

**DOI:** 10.1101/2024.03.26.586761

**Authors:** Imtiaz Ahmad, Kis András, Radhika Verma, István Szádeczky-Kardoss, Henrik Mihály Szaker, Aladár Pettkó-Szandtner, Dániel Silhavy, Zoltán Havelda, Tibor Csorba

## Abstract

Regulation of transcriptional machinery and its adaptive role under different stress conditions are studied extensively in the dicot model plant *Arabidopsis*, but our knowledge on monocot species remains elusive. TFIIS is an RNA polymerase II associated transcription elongation co-factor. Previously it was shown that TFIIS ensures efficient transcription elongation that is necessary for heat stress survival in *A. thaliana*. However, the function of TFIIS has not been analysed in monocots. In the present work, we have generated and studied independent *tfIIscrispr* mutant barley lines. We show that, TFIIS is needed for reproductive development and heat stress survival in barley. The molecular basis of HS-sensitivity of *tfIIs* mutants is the retarded expression of heat stress protein transcripts, that leads to late accumulation of HSP chaperones, enhanced proteotoxicity and ultimately to lethality. We also show that TFIIS is induced and autoregulated in response to heat, supporting a conserved adaptive function of these control elements for plant thermal adaptation. In sum, our results are a step forward for the better understanding of transcriptional machinery regulation in monocot crops.

## Introduction

Monocot species are important agronomic plants. Their grains contribute enormously to global food supply and animal nutrition. Despite the rising demands, crop plants are understudied. To increase their productivity the basal understanding of development and stress resilience is of uttermost importance.

Barley (*Hordeum vulgare*) ranks fourth in terms of production, following wheat, maize, and rice. Worldwide barley grain production reached above 150 M tonnes in 2022/2023 vegetation period(http://faostat3.fao.org)(https://www.statista.com/statistics/271973/world-barley-production-since-2008/#:∼:text=The%20global%20production%20volume%20of,metric%20tons%20in%202021%2F2022.). Barley grain is used as an animal feed, for production of different beverages and in human nutrition, as it has high nutritional value for its starch, protein, fibre, and micronutrient contents. Barley has emerged as a monocot crop research model lately (Harwood, 2019). There are numerous benefits of using barley to dissect diverse molecular and physiological pathways. It is a self-pollinating plant, has a diploid genome which was recently published, a wealth of genetic variants and ecotypes are available, it grows in a wide range of environmental conditions and climates and is easy to handle in laboratory (Harwood, 2019; Jayakodi *et al*., 2020).

Global warming and climate change endanger survival of natural habitats worldwide, also causing great losses in productivity of crops (Akerfelt *et al*., 2010; Yeh *et al*., 2012; Ohama *et al*., 2017). Extreme heat is detrimental to cells and organisms, as it causes disorganisation of lipid membranes, induces damage to nucleic acids, and leads to protein misfolding and aggregate formation. There are several mechanisms that work to diminish the adverse effects of high temperature; these are collectively called heat stress response (HSR). Transcriptional regulation is a central element of HSR. Master regulators of the HSR are the Heat Stress Factors (HSF). Upon HS the HSFs are activated, they bind to heat shock elements (HSE), *cis* regulatory motifs of HS-responsive loci, and switch on/off multiple transcriptional cascades causing the production of secondary HSFs, heat stress protein (HSP) chaperones, and antioxidant enzymes among others. HSPs and antioxidant factors re-establish cellular proteostasis and reorganise physiology to optimise energy usage (Akerfelt *et al*., 2010; Yeh *et al*., 2012; Ohama *et al*., 2017). The timely and qualitative transcriptional reprogramming is essential for an efficient HSR (Akerfelt *et al*., 2010; Ohama *et al*., 2017; Szadeczky-Kardoss *et al*., 2022; Obermeyer *et al*., 2023).

RNA transcription is a central act of cellular life that enables the flow of genetic information from the genome to orchestrate cellular pathways. The constant alterations of transcription lays at the basis of development and provide an adaptation capacity to the persistently changing environment. In eukaryotes, the RNA Polymerase II (RNAPII) complex is responsible for the production of protein-coding and many non-coding transcripts. It was shown in *A. thaliana* that upon encountering heat stress a massive part of transcription is shut-down (mostly genes acting in development), while at the same time the stress-responsive genes are hyper-transcribed (Akerfelt *et al*., 2010; Yeh *et al*., 2012; Ohama *et al*., 2017; Szadeczky-Kardoss *et al*., 2022). Mechanisms leading to increased transcriptional output include the elevated level of transcriptional initiation and/or a faster elongation rate. While transcriptional initiation control was studied thoroughly, much less is known about transcriptional elongation control. Gathering evidence suggests that the precise regulation of transcriptional elongation is needed for the adequate response to environmental alterations (Antosz & Deforges, 2020; Szadeczky-Kardoss *et al*., 2022; Obermeyer *et al*., 2023).

Transcriptional elongation is irregular, proceeding with variable speed and efficiency. Several transcription elongation factors including TFIIS, PAF1c, FACT, DSIF, SPT6 etc. assist the RNA polymerase II (RNAPII) during the transcriptional process (Antosz *et al*., 2017; Grasser & Grasser, 2018; Aoi & Shilatifard, 2023). Transcription may be interrupted by DNA damages, transcriptional mistakes, or roadblocks. In such cases the RNAPII complex is arrested and may backtrack on the template. During backtracking the 3’ terminus of the nascent RNA is displaced from the RNAPII catalytic core. To resume transcription, the protruding RNA is cleaved off by the intrinsic exonuclease activity of RNAPII. The cleavage reaction by RNAPII in itself is slow, but it’s greatly accelerated by the elongation factor TFIIS (Kettenberger *et al*., 2003; Sydow & Cramer, 2009; Ruan *et al*., 2011; Xu *et al*., 2017).

TFIIS protein is composed of three functional domains: the N-terminal domain I involved in nuclear localization and serves as a scaffold for protein co-factor interactions (Cermakova *et al*., 2021; Cermakova *et al*., 2023); the central domain II facilitate interaction with RNAPII core complex (Kettenberger *et al*., 2003; Xu *et al*., 2017); and domain III helps to perform the nucleolytic catalysis. Domain III consists of three **β**-sheets that are held together by four cysteine residues chelating a zinc ion. Following RNAPII arrest, the Zn-finger domain is inserted through a pore, reaching adjacent to the RNAPII polymerization/exonuclease core site. At the tip of the domain there is a highly conserved dipeptide (aspartic acid, glutamic acid, DE) motif that complements the active site of RNAPII by stabilisation of the loosely bound Mg^2+^ needed for the nucleolytic reaction. The presence of the DE dipeptide accelerates RNA cleavage and quickly reinstalls productive transcription.

The activity of TFIIS is vital for heat stress tolerance in *A. thaliana* (Szadeczky-Kardoss *et al*., 2022; Obermeyer *et al*., 2023). Based on its HS-inducibility we proposed that TFIIS role in stress adaptation is conserved in plants. In the present work we study the biological roles of TFIIS in barley. Our data indicate that in monocots, TFIIS is needed for normal reproductive development at ambient temperature and for survival during heat stress.

## Materials and methods

### Plant materials, growth conditions and heat stress sensitivity treatments

Barley (*H. vulgare* L. cv. ‘Golden Promise’, wild type, wt) plants were grown in light cabinets (MLR-350; Sanyo, Tokyo, Japan) at 20 °C, long day (LD, 16 h light/ 8 h dark) conditions (Hamar *et al*., 2020). For the heat stress (HS) treatments, wild-type and mutant plants were pre-grown for a week in Jiffys then potted side-by-side, one wt plant besides a mutant plant in the same pot and were grown for an additional week. The 14 days old plants were subjected to persistent moderate heat stress (Thermotolerance to Moderately High Temperature regime, TMHT) consisting of 40 °C treatment for 1-3 days. After treatment the plants were placed back to ambient conditions (20 °C, LD) and recovered for an additional 14 days. HS-sensitivity documentation was performed on day 30. For molecular works (RNA and protein content examination), leaf samples were collected from control and treated plants immediately following the treatments such as: non-treated (NT), one hour at 40 °C (1 h), two hours at 40 °C (2 h) and one day (1 d), frozen in liquid nitrogen and stored at −70 °C until use.

### Generation of CRISPR mutants

The barley TFIIS locus and genomic sequence was identified previously using the Ensembl Plants database (*HORVU*.*MOREX*.*r3*.*5HG0524690*, formerly *HORVU5Hr1G111700* (Szadeczky-Kardoss *et al*., 2022)). To design CRISPR cleavage target sites we employed the CRISPOR software (Concordet & Haeussler, 2018). sgRNAs exhibiting minimal off-target activities were selected for generation of two mutant classes: sgRNA1 targets the gene locus in the first exon of the protein, creating the loss of the complete protein while the sgRNA2 induces mutations just upstream of the catalytically active TFIIS-type Zn-finger domain. The CRISPR-Cas9 vector containing the sgRNAs was prepared as described previously (Kis *et al*., 2024).

For mutant plant generation immature barley embryos were transformed using *Agrobacterium* AGL1 strain. Transgenic plants were generated from agrobacterium-infected calli. Genomic DNA was extracted from each plant using Extraction and Dilution buffers (Sigma-Aldrich, E7526 and D5688, respectively, www.sigma-aldrich.com). Target sites were amplified using Phusion High-Fidelity DNA Polymerase (Thermo Fisher Scientific, F537S) from wt and transgenic lines. The presence of mutation was identified by T7 Endonuclease digestion (NEB M0302, www.neb.com). Homozygous lines were obtained for further study. Plants were grown in greenhouse conditions.

Primers used for cloning and target site amplification are listed in the *Supplementary Materials*.

### Embryo rescue

Fully mature barley seeds of wild type and *tfIIs-cr1* mutant line were immersed in water and incubated in dark for one day. Subsequently, embryos were extracted from seeds using forceps and the embryos plated with scutellum side-up on callus induction medium containing 4.3 g l^-1^ Murashige & Skoog plant salt base (Duchefa, www.duchefa.com), 30 g l^-1^ Maltose, 1.0 g l^-1^ Casein hydrolysate, 350 mg l^-1^ Myo-inositol, 690 mg l^-1^ Proline, 1.0 mg l^-1^ Thiamine HCl, 2.5 mg l^-1^ Dicamba (Sigma-Aldrich, www.sigma-aldrich.com) and 3.5 g l^-1^ Phytagel (Bartlett *et al*., 2008). After roots and shoots have emerged, seedlings were transferred to Jiffys and further grown in light cabinets for subsequent experiments or heat stress treatments.

### RNA extraction and qRT-PCR

RNA extraction from 30-60 mg barley plant leaves was performed by the phenol:chloroform method. For qRT-PCR assays, 2,5 μg total RNA was DNase treated according to manufacturer’s instructions (NEB, M0303, www.neb.com), precipitated in ethanol, resuspended in sterile water. 1 μg of DNase-treated RNA was used for the first-strand complementary DNA reaction with random primers, according to the manufacturer’s instructions (NEB, E6300, www.neb.com). qPCRs were done using qPCR Master Mix (NEB, M3003, www.neb.com) according to the manufacturer’s instructions. qPCR reactions were run in a Light Cycler 96 (Roche) Real-Time PCR machine. At least three biological replicas were assessed in each experiment and standard error bars displayed. P values were calculated using unpaired two-tailed Student t-test to assess the significance of differences. For DNA oligonucleotides used in the study please see the *Supplementary Materials*.

### Western blot analysis

Barley leaf (middle section of the second leaf) samples were homogenised in 2x SDS-PAGE buffer (100 mM Tris–HCl, pH 6.8, 20% glycerol, 2% SDS, 2mM DTT, 0,05% Bromophenol Blue), boiled for 5 min and cell debris removed by centrifugation at 14 000 x g at 4 °C for 10 min. The supernatants were resolved on 10% SDS-PAGE, transferred to Hybond PVDF membranes (GE Healthcare) and subjected to western blot analysis. Antibodies used for detection: anti-HSP70 antibody (AS08 371, Agrisera), anti-HSP90 antibody (AS08 346, Agrisera), anti-HSP101 (AS07 253, Agrisera); as secondary antibody, we used monoclonal HRP-conjugated anti-rabbit (A6154, Sigma-Aldrich) antibody. The proteins were visualised by chemiluminescence (Clarity ECL reagent; Bio-Rad, www.bio-rad.com) and quantified by Image Lab 5.1 (Bio-Rad). Rubisco large subunit (RbcL) stain free signals were used as loading control.

### Protein aggregate purification

Protein aggregate detection was done as described before (Szadeczky-Kardoss *et al*., 2022). Protein extracts were prepared by homogenization of 0.1g plant leaf fresh weight material in 2.4 ml of isolation buffer [25 mM HEPES, pH 7.5, 200 mM NaCl, 0.5 mM Na2EDTA, 0.1% (v/v) Triton X-100, 5 mM ε-amino-N-caproic acid, 1 mM benzamidine], using a mortar and pestle and then a Cole-Parmer PTFE glass tissue grinder. The soluble and insoluble fractions were separated from 2 ml of total extract by centrifugation at 16 000 x g for 15 min at 4 °C. The soluble fraction was denatured by adding 0.5 volume of 2x SDS-PAGE buffer and heating for 5 min at 95 °C. The insoluble pellet was washed five times repeatedly by resuspension in the isolation buffer containing 0.1 g of quartz sand (Sigma-Aldrich) and vortex. The insoluble pellet was resuspended in 400 ml 2× SDS-PAGE sample buffer and clarified by centrifugation at 1500 × g for 1 min (insoluble fraction). Samples were separated by SDS-PAGE and stained with Coomassie Blue and later Silver Staining method. The whole lanes of insoluble fractions have been quantified by Image Lab 5.1 (Bio-Rad) and ratios to Rubisco large subunit (RbcL) stain free signals was calculated.

### Bioinformatic tools and analysis

To analyse sequence conservation of TFIIS homologues, sequences were identified and extracted from the Ensamble Plants and Uniprot (Martin *et al*., 2023; UniProt, 2023) databases. Protein alignments were made using Jalview software (www.Jalview.org) (Waterhouse *et al*., 2009). The prediction and illustration of the 3D structure of TFIIS domain II were acquired from and generated by AlphaFold DB (Jumper *et al*., 2021).

## Results

### TFIIS conservation in monocots

To assay the importance of TFIIS roles in cereals, first we analysed protein sequence conservation. For this, we identified protein sequences of TFIIS homologues in main monocotyledonous crops, and a few representative species including gymnosperms, early dicotyledonous and eudicotyledonous plants, that were used for comparison, and performed alignments (Fig S1, S2). These revealed that the three domains of TFIIS protein homologues are highly conserved in all studied plant groups (Fig 1), including the 5 α-helices of domain I (Fig S1A), the 7 α-helices of domain II (Fig S2A), and the 3 ***β***-sheets of domain III (Fig S1B). The 4 cysteine residues and DE dipeptide of domain III zinc finger are also present in all lineages (Fig S1B), suggesting that these TFIIS homologues may be indeed functional. Besides the few scattered amino acids and some lineage-specific differences, we observed a monocot-specific stretch of amino acid insertion in domain II between the *α6* and *α7*-helices; we named these *α6-b* and *α6-c*. The insertion consists primarily of charged amino acids (Fig S2A). Based on the *in-silico* prediction of *Arabidopsis* and maize (*Zea mays*) TFIIS protein structures (Fig S2B), the *α6-b* and *α6-c* helices are located on the outer surface of RNAPII-bound TFIIS.

**Figure 1:**
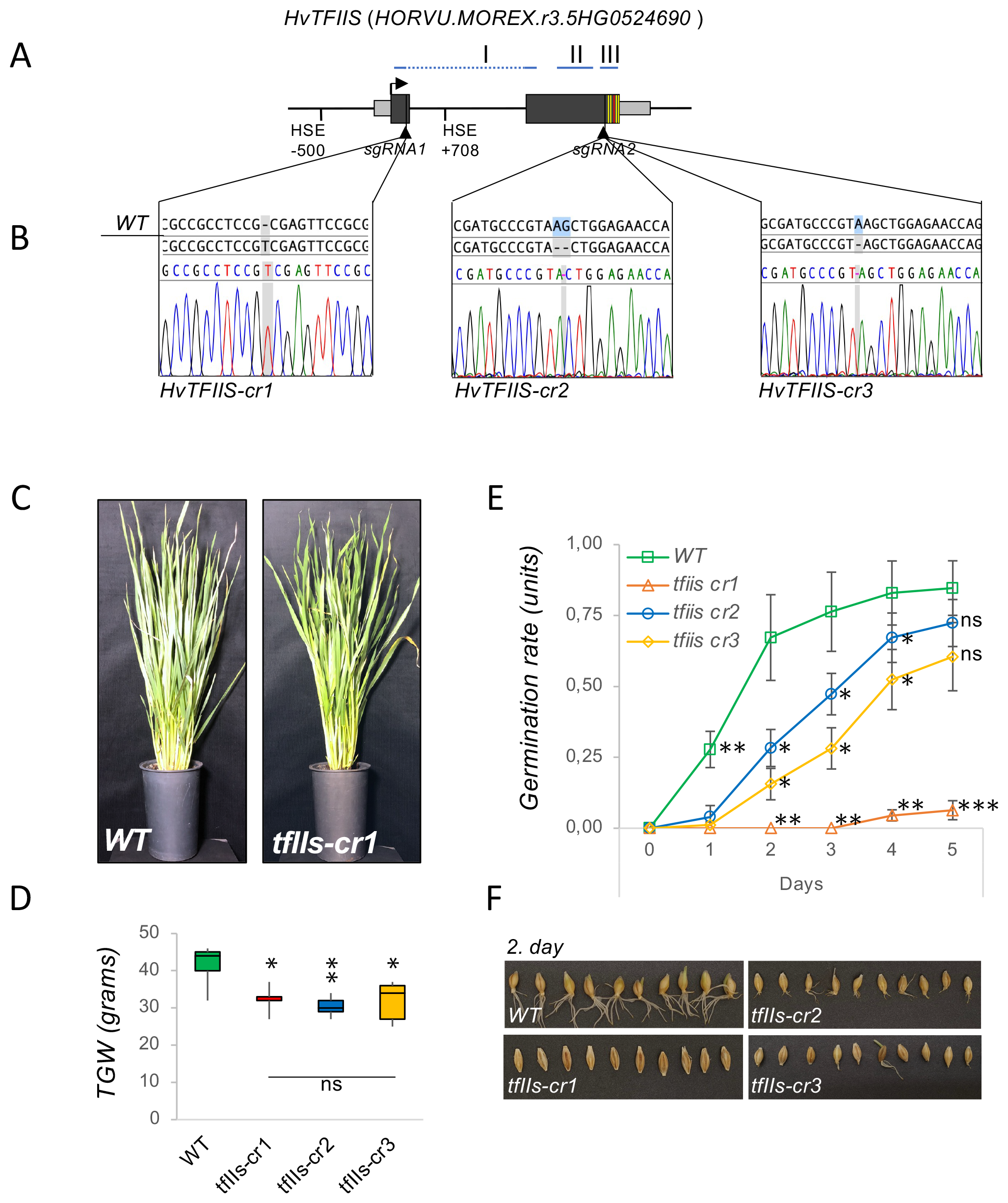
TFIIS is needed for reproductive development in barley. (A) Schematic representation of *HvTFIIS* gene locus; protein domain I, II and III location is shown above; exons as black boxes, UTRs as grey boxes, Zn finger domain cysteine residues and DE acidic dipeptide motif as yellow and red line, respectively; HSE *cis* elements and location of sgRNA guide sites are shown below; (B) TFIIS mutants of barley were created by CRISPR mutagenesis; insertion or deletion mutation within the TFIIS locus in the selected transgenic lines are shown on chromatograms; (C) *tfIIs-cr1* mutant plants have slightly retarded vegetative growth compared to wild type plants; (D) The calculated thousand grain weight (TGW) of wild type and *tfIIs-cr* mutant plants; (E) Germination rate quantification of wild type and *tfIIs-cr* mutant plants during a timeseries; (F) Pictures of germinating seeds on 2. day; bars represent standard errors based on at least three bio reps; P-values based on two-tailed Student’s t-test (*P < 0.05, **P < 0.01, ***P < 0.001, ns; non-significant) show differences between wild type and mutant plants.

### Generation of independent lines of *TFIIS*-mutant barley

To uncover the function of TFIIS during development and stress adaptation in monocot crops, we decided to create independent TFIIS mutant lines in barley; for this we used CRISPR-mutagenesis (*see Materials and Methods*). Two different sgRNA target sites were chosen for the work: sgRNA1 targets the TFIIS locus within the first exon of the gene; the sgRNA2 targets the locus upstream to the catalytic active Zn-finger domain (Fig 1A, B and S3A, B). We selected two independent lines derived from each sgRNA-targeted progeny and obtained homozygous mutants. Each of these lines harboured a frame-shift mutation, such as an adenine insertion and a thymine insertion, in the CRISPR/sgRNA1 lines, and an adenine-guanine (AG) dinucleotide deletion or an adenine deletion, in the case of CRISPR/sgRNA2 lines, respectively (Fig 1B).

The sgRNA1-targeted mutagenesis induced frameshift mutations that resulted in complete loss of the TFIIS protein (knock-out mutant, Fig S3B). As it turned out later, one of these homozygous lines (+A) was sterile therefore could not be used for further work (*not shown*). The other sgRNA1-induced insertion line (+T, that was named *tfIIs-cr1*) could be propagated by embryo rescue (*see later*). We failed to find any other sgRNA1-generated *tfIIs* mutant of several tested able to produce viable homozygous progenies. The sgRNA2-directed mutations resulted in frameshifts at the end of the gene; these alleles may produce truncated protein bearing the intact N-terminal and the central domain but lacking the entire Zn-finger of catalytic domain III (antiarrest activity mutant, Fig S3B). The two independent homozygous lines generated using the sgRNA2 were both fertile and produced viable seeds. These lines were named *tfIIs-cr2* and *tfIIs-cr3*, respectively. The three independent lines *tfIIs-cr1, tfIIs-cr2* and *tfIIs-cr3* alongside wild type *Golden promise* (wt) were used for further experimentations.

### HvTFIIS is needed for efficient reproductive development

First, we inspected the development of *tfIIs* mutants under ambient conditions (Fig 1C-F, S3C). The *tfIIs-cr1* plants were somewhat retarded in their growth compared to wt (Fig 1C). The growth of *tfIIs-cr2* and *tfIIs-cr3* plants was very similar to wt plants in all aspects of vegetative development including leaf shape and overall plant stature (Fig S3C). Reproductive development, however, was affected in all *tfIIs* mutants. The *tfIIs-cr1* seeds were slightly elongated and unable to germinate, even when pre-treated in dark with gibberellic acid to promote root and shoot development (Fig 1E, F). The germination incompetent phenotype was consistent on multiple plants in consecutive cultivation periods. We have rescued the embryos by extracting them from the mature seeds; when placed on growth media (Fig S3D, *see also Materials and Methods*), they could develop roots and shoots, and finally progressed into mature plants. Wild type plant embryos were handled in the same way to produce control material for HS-sensitivity experiments or molecular analysis. Reproductive development was also affected in the *tfIIs-cr2* and *tfIIs-cr3* plants although to a lesser extent: the seeds of both lines had a significantly retarded germination rate (Fig 1E, F). Besides the reduced germination capacity, the *tfIIs-cr1, -cr2* and *-cr3* lines had a thousand grain weight (TGW) reduced to 75%, 68% and 77%, respectively, compared to wt (Fig 1D). These data combined suggest that TFIIS is needed under ambient conditions for ensuring the reproductive fitness of barley plants.

### TFIIS is a basic factor of efficient HSR

Previously it was shown that TFIIS is needed for survival of plants exposed to heat stress (Szadeczky-Kardoss *et al*., 2022; Obermeyer *et al*., 2023). To analyse the requirement of TFIIS in barley high temperature adaptation, we subjected the *tfIIs* mutant lines to Thermotolerance to Moderately High Temperature (TMHT) stress regime. All *tfIIs-cr* plants were heat-sensitive, showing mild symptoms after 1d TMHT (40°C) and strong retardation or lethality after 2d/40°C treatment (Fig 2A, B and S4A, B).

**Figure 2:**
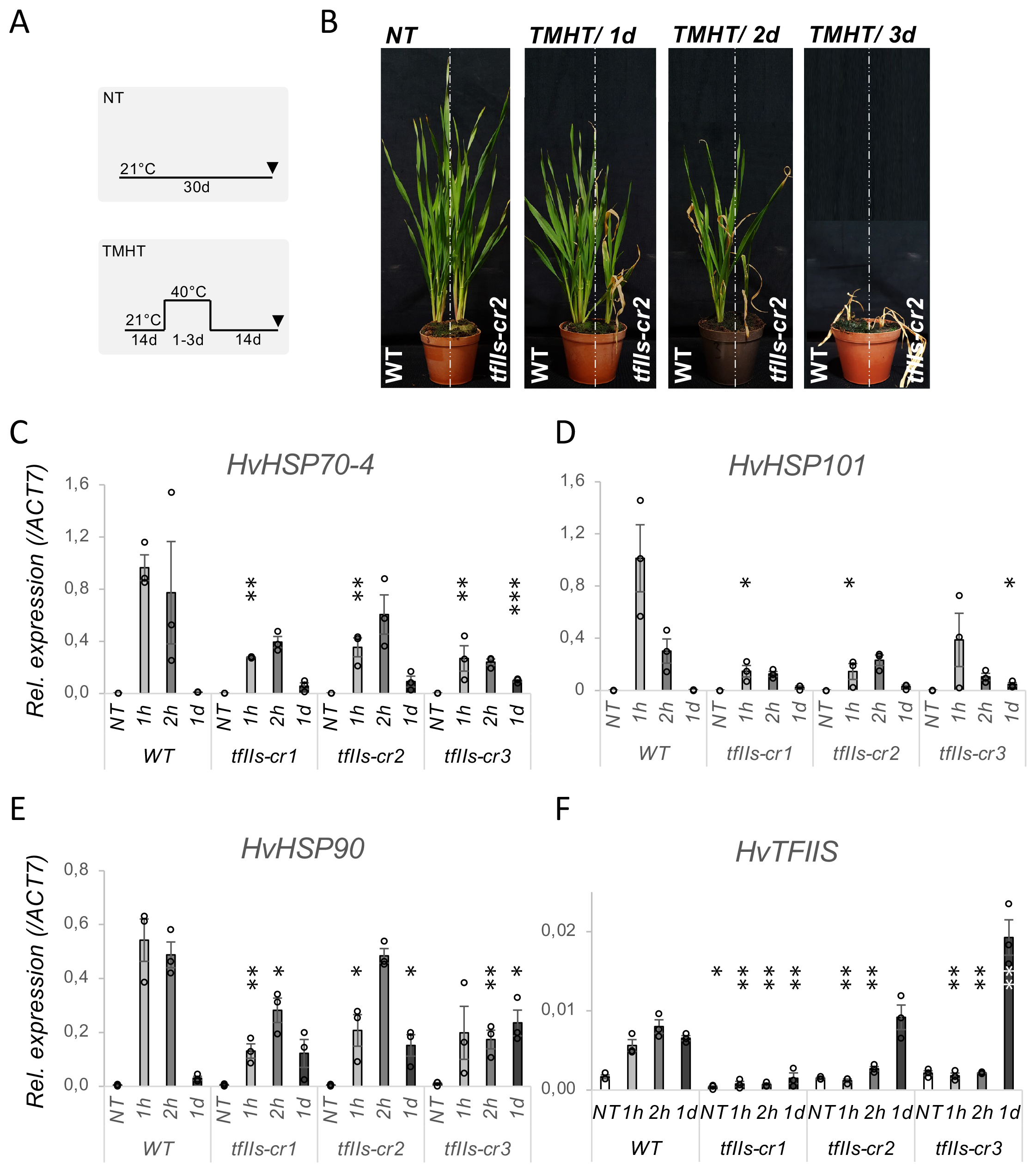
TFIIS is indispensable for efficient heat stress response. (A) Schematic representation of non-treated (NT) and Thermotolerance to Moderately High Temperature regime (TMHT); arrowheads show the time of sampling; (B) wild-type (wt) and *tfIIs* CRISPR mutants (*tfIIs-cr*) were exposed to NT or TMHT for 1 hour (1h), 2h or 1 day (1d); Accumulation of HSR-transcripts is retarded altered in the absence of HvTFIIS. Relative expressions of (C) *HSP70-4*, (D) *HSP90* and (E) *HSP101* and (F) *TFIIS* mRNA during a timeseries; bars represent standard errors based on three bio reps; P-values based on two-tailed Student’s t-test (*P < 0.05, **P < 0.01, ***P < 0.001) show differences between wild type and mutant plants.

To decipher molecular alterations at the basis of TFIIS actions in barley during HSR, we analysed mRNA changes of representative HSP transcripts during a time series. Accumulation of HSP transcripts *HSP70-4, HSP90* and *HSP101* in response to early heat exposure (1-2h/40°C) was significantly retarded in *tfIIs-cr* mutant plants compared to wt (Fig 2C-E). Upon persistent heat stress (1d/40°C), in the wt plants expression of these HSPs was efficiently attenuated; however, in the *tfIIs* mutant plants remained elevated.

To investigate whether the mRNA changes have a perceivable outcome at protein level, we quantified protein amount changes: HSP70, HSP90 and HSP101 accumulated efficiently in the wt plants but to a significantly lower level in *tfIIs-cr* mutants, especially observed in the early HS period (1-2h/40°C)(Fig 3A-C, S4C). In the late HS most differences were moderated, suggesting that in the long term the *tfIIs* mutant plants can partially offset the early inequality in their HSP chaperon contents, likely by extending the period of mRNA transcription and/or protein translation.

**Figure 3:**
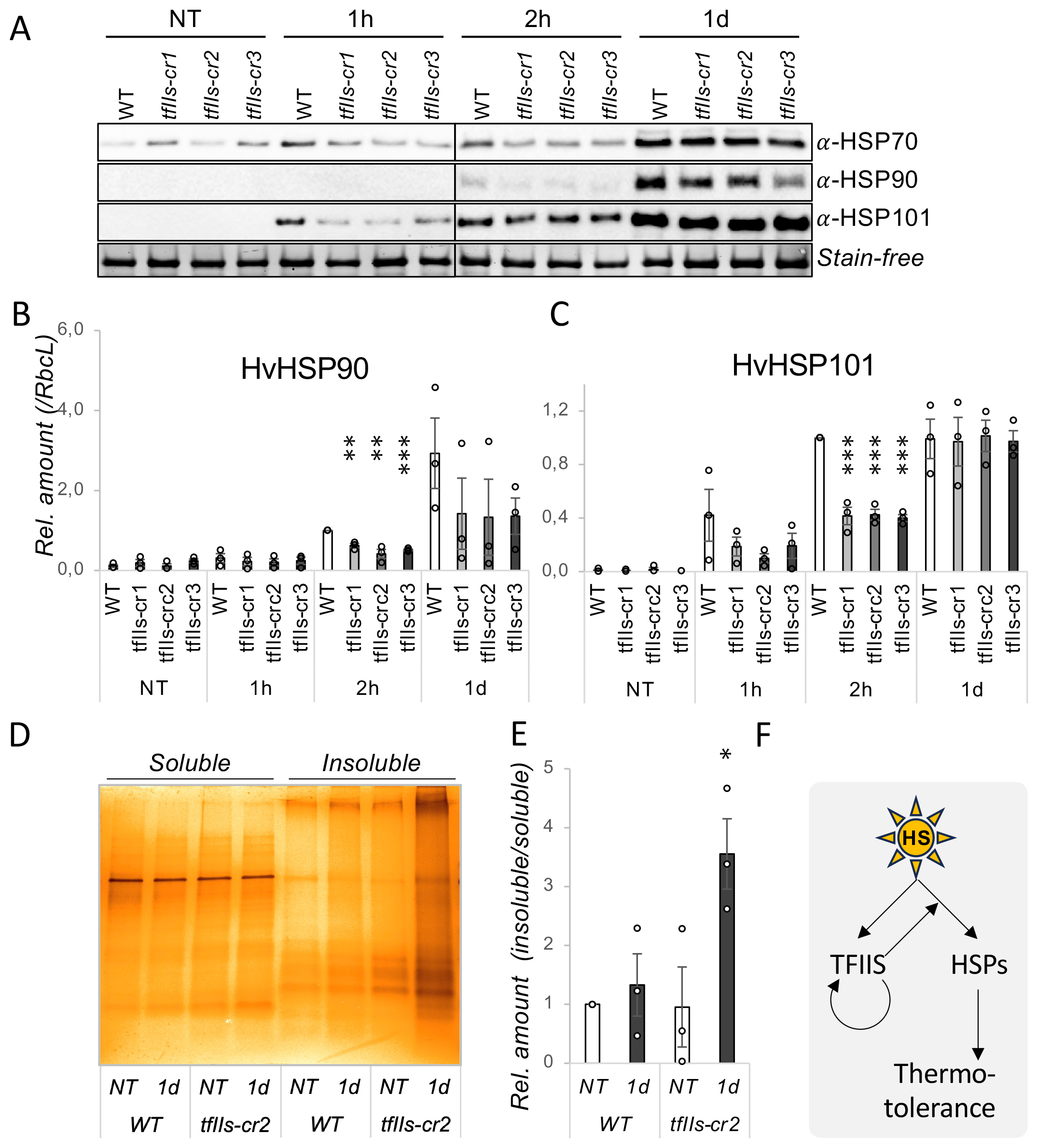
Heat stress causes enhance proteotoxic stress in absence of TFIIS. (A) Western blot images HSP70, HSP90 and HSP101 proteins; stain-free picture is shown as a loading control; (B, C) Quantification of protein accumulation of HSP90 (B), and HSP101 (C) during the same timeseries; wild-type (wt) and *tfIIs* CRISPR mutants (*tfIIs-cr1, -cr2 and -cr3*, respectively) were exposed to NT or TMHT for 1 hour (1h), 2h or 1 day (1d), as shown; (D) Silver staining gel images of insoluble protein fractions from wt and *tfIIs-cr2* plants exposed to non-treated or 1d TMHT heat treatment; Rubisco large subunit (RbcL) is shown as loading control of soluble fraction; treatment conditions as in Fig 2. (E) Ratio quantifications of insoluble/soluble protein fractions in wild type and *tfIIs-cr2* mutant plants at non-treated (NT) and TMHT/1d (1d) stress conditions; P-values based on two-tailed Student’s t-test (*P < 0.05, **P < 0.01, ***P < 0.001) show differences between wild type and mutant plants. (F) Working model for TFIIS action during heat stress response: elevated temperature induces expression of HSPs and TFIIS; TFIIS elongation factor positively feed-back regulates itself and further enhances expression of HSP transcription; the timely accumulation of HS-transcripts and proteins ensures efficient thermotolerance in barley plants.

### Retarded HSP production in the absence of TFIIS causes proteotoxicity

To assess the downstream consequences of under-accumulation of HSP chaperones involved in protein homeostasis including refolding of denatured proteins, disaggregation of protein precipitates and promotion of decay of irreversibly unfolded proteins, we analysed protein aggregate levels. For this, we purified insoluble protein fraction and calculated insoluble/soluble protein ratios in non-treated and 1d heat-treated wt and *tfIIs-cr2* plants. Significantly higher accumulation of insoluble proteins was observed in the *tfIIs-cr2* plants in response to heat stress, suggesting a proteotoxic condition (Fig 3D, E).

### TFIIS is HS- and self-regulated

To better understand how TFIIS itself is regulated during HSR, we investigated its mRNA dynamics in response to heat. *TFIIS* mRNA level increased with 3- or 5-fold factor, respectively, in the 1h or 2h/40°C early HS treatment (Fig 2F). Besides, the *TFIIS* transcript level remained elevated at 1d, suggesting a need for sustained TFIIS activity. *HvTFIIS* locus contains two canonical HSE *cis* elements (Fig 1A); these are likely involved in HS-induced transcriptional regulation of the locus.

In the *tfIIs-cr1* mutant the level of TFIIS mRNA was significantly lower under NT conditions and has shown only a marginal (non-significant) accumulation trend in response to heat. The single nucleotide insertion (+T) in this line causes frameshift within the coding region of TFIIS locus, leading to the appearance of a premature termination codon (PTC)(Fig 1 and S3). Because of this, the *tfIIs-cr1* mRNA has an intron containing, long 3’UTR. The long 3’UTR and introns within the UTR are the two main canonical features recognized by the Nonsense mediated decay (NMD) RNA quality control pathway (Kerenyi *et al*., 2008; Kurosaki *et al*., 2019; Ohtani & Wachter, 2019). The low levels of *TFIIS* transcript in *tfIIs-cr1* and its inability to accumulate in response to HS are very likely the consequence of NMD-mediated decay.

In the *tfIIs-cr2* and *tfIIs-cr3* lines the *TFIIS* mRNA level under ambient conditions was similar to wt. In the early heat response (1-2h/40°C), contrarily to wt, there was no accumulation of *TFIIS* mRNA (Fig 2F). This implies that rapid induction of *TFIIS* mRNA expression directly or indirectly requires the activity of TFIIS protein itself. Oppositely to the expression pattern observed during early heat stress response, under persistent HS (1d/40°C), a strong accumulation of *TFIIS* transcripts was noticed. This induction at a later stage of HSR suggests the presence of a secondary, TFIIS independent regulatory mechanism that may serve as a feed-back compensation to replenish TFIIS.

Taken together, the data points to the direction that TFIIS is much needed for early and late HSR, and that separate mechanisms operate to ensure optimal expression of TFIIS under early and late HSR.

## Discussion

Cereal crops are cultivated worldwide since their grains constitute the basis of animal and human nutrition. Deciphering the molecular mechanisms that coordinate development and stress response pathways in crops is of great interest. Although transcriptional regulation is central for transferring genetic information to coordinate cellular machinery, the available knowledge in monocot species is very limited. Elongation factors play a crucial role in transcriptional regulation, as well documented in dicots and non-plant systems (Durr *et al*., 2014; Antosz *et al*., 2017; Conaway & Conaway, 2019; Godoy Herz *et al*., 2019; Cermakova *et al*., 2021; Michl-Holzinger *et al*., 2022; Obermeyer *et al*., 2022; Szadeczky-Kardoss *et al*., 2022; Aoi & Shilatifard, 2023; Obermeyer *et al*., 2023). Research on transcriptional elongation control in monocots is scarce. It was shown that the DSIF complex subunits SPT4 and SPT5 paralogs are needed for proper vegetative and reproductive growth in rice (Liu *et al*., 2023). A hypomorph *spt5-1* mutant rice displays semi-dwarfism and has a shortened life cycle. Furthermore, *spt4* or *spt5-1* homozygous mutant rice progenies could not be obtained, while heterozygous plants had impaired grain development (Liu *et al*., 2023). A genome-wide association study identified Transcription Elongation Factor 1 (TEF1) (10.1016/j.jia.2024.03.030) to be involved in salt stress tolerance in *Sorghum*, underpinning the critical role of transcriptional elongation control in regulation of abiotic stress responses. To better understand the importance of transcriptional elongation regulation in cereals, we focused our research on TFIIS elongation factor in the barley model. We created independent CRISPR lines using two different sgRNAs as guides and have selected independent transformants to exclude phenotypic alterations derived from off-target editing or transgene insertional events (Fig 1, S3). We observed a marginal reduction in the vegetative growth of *tfIIs-cr1*, but no obvious developmental alterations in the other two independent lines, *tfIIs-cr2* and *-cr3* (Fig S3C). The experiments were performed on multiple plants within each line, for two consecutive vegetation periods. The limited alterations of TFIIS’s absence on growth are in concert with data obtained from *Arabidopsis*, where the *tfIIs* mutants display normal phenotype (Grasser *et al*., 2009). These observations suggest limited impact of TFIIS on vegetative development under ambient conditions.

However, in barley, unlike in Arabidopsis, the absence of TFIIS, caused severe reproductive fitness cost at ambient temperature, as revealed by qualitative and quantitative features of the grains: (i) homozygous progenies from sgRNA1 transformants were obtained from only two lines and only one of them was fertile (the *tfIIs-cr1*), (ii) the *tfIIs-cr1* seeds are germination incompetent, (iii) the *tfIIs-cr2* and *tfIIs-cr3* seeds’ germination capacity is significantly decreased, and (iv) the TGW of all *tfIIs-cr* transgenics is significantly reduced to about three-quarter compared to wt plants and non-significantly different between each other (Fig 1D-F). These are robust findings and are in close unison with data obtained in *osstp4* and *osstp5-1* rice mutants (Liu *et al*., 2023). Although reproductive development was impaired in all three *TFFIIS* mutants, the differences of germination phenotypes of *tfIIs-cr1* (not germinating) and *tfIIs-cr2* and *-cr3* seeds (having retarded germination) are obvious. The *tfIIs-cr1* is very likely a null-mutant, the mRNA is barely detectable, it is not inducible and if it is translated, the generated protein would lack all three domains of TFIIS protein. By contrast, the *tfIIs-cr2* and *-cr3* mRNAs are expressed and inducible; likely they encode truncated proteins containing domain I and II and a non-functional domain III. TFIIS has dual role in transcriptional regulation: domain I act as a scaffold of the transcriptional machinery by tethering elongation cofactors (Cermakova *et al*., 2021) while domain III stimulates the exonucleolytic activity of PolII (Kettenberger *et al*., 2003). We propose that in the *tfIIs-cr2* and *-cr3* antiarrest mutants the milder reproductive phenotype could be due to the retained scaffold/protein interactor and/or Pol II interaction function. A less likely alternative is that the sgRNA1 guide has a special off-target, that enhances the reproductive phenotype of *TFIIS* mutation. Further work is necessary to map the different putative functions linked to the specific domains of TFIIS. Previously it was shown that the efficient transcriptional reprogramming is vital for temperature adaptation in *A. thaliana* (Szadeczky-Kardoss *et al*., 2022; Obermeyer *et al*., 2023). Here we extend findings in *Arabidopsis* and show that TFIIS is needed for efficient HSR and stress survival in barley. As molecular changes and the HS-sensitivity phenotype was consistent for both mutant groups (the knock-out *tfIIs-cr1* and the antiarrest *-cr2* and *-cr3* mutants) tested, we postulate that the transcriptional antiarrest activity of TFIIS is a requirement for efficient HSR in barley. In support of these, we have shown that TFIIS is induced by HS and may be autoregulated. Autoregulation of TFIIS was also verified in *Arabidopsis*, suggesting that it is an important element of transcriptional control in plant kingdom, at least under HS conditions. Our observations further suggest that transcription elongation efficiency in *tfIIs* mutant barley may be diminished, and transcriptional program switch could not occur at the required pace and quality which leads to proteotoxicity and finally lethality (Fig 3F). A transcriptome analysis would uncover the full spectrum of differentially expressed transcripts during HSR and map transcriptional regulatory functions linked to the different domains of TFIIS by comparison of the two *tfIIs* mutant groups.

Lastly, we analysed protein sequence conservation of TFIIS homologs in monocot species by comparing them to homologs in gymnosperms, early dicots, and eudicot plants. Strong conservation of TFIIS protein sequence and secondary motifs including α-helices, β-sheets, and the Zn-finger domain features was evidenced. We have noticed a sequence expansion consisting mainly of acidic residues in the region between *α6-α7* on domain II in plant kingdom ((the *α6-α7* region is short in yeast, worm, fly or mammalian, *data not shown*, (Kettenberger *et al*., 2003)), suggesting plant-specific roles. This amino acid stretch is further expanded and *in silico* predicted to form additional α-helix structures in monocot species (*α6-b, α6-c*). Whether the charged surface of *α6-b/α6-c* (together with *α7*) helices (Fig S2B) aids better protein solubility, prevention of aggregation or interactions with protein partners, remains to be determined. Notably, *Brassicaceae* plants (including *A. thaliana*) have an aromatic tyrosine residue emerging from the acidic surface of this area. This constitutes a strong hydrophobic point that may serve as a binding “handle” for protein interaction partners. Proteins with binding pocket able to accommodate the tyrosine aromatic ring have been described before (Wonderlich *et al*., 2008); interestingly tyrosine-dependent recruitment of cofactor protein is further stabilised by acidic residues in proximity (Wonderlich *et al*., 2008). In sum, these observations hint to the possibility that TFIIS *α6-α7* on domain II may be a platform for client protein binding through charge-charge interactions in plants. We speculate that cofactor binding may be further enhanced by surface expansion (e.g. *α6-b/α6-c* helices) in monocots or by the Tyr-binding in *Brassicaceae*. Future work may unravel protein partners tethered by TFIIS to the core RNAPII to modulate its functions.

In sum, our data prove the role of TFIIS in reproductive development and high temperature adaptation of barley. The results endorse previous findings in *Arabidopsis* and certify conservation of TFIIS roles (notably, *Arabidopsis* and barley are considerably diverged, their separation was estimated to be 140 Mya (Moore *et al*., 2007)). Further systemic study on transcriptional regulation could uncover commonalities or peculiarities of the pathway within monocot species. This knowledge combined with the now available genomic information may help directional selection and breeding programs to increase productivity and resilience of these important plant species.

## Supporting information

Supplementary figures and tables supporting the work.

## Supplementary information

The online version contains supplementary material available at https://doi.org/…

## Author contribution statement

IA, AK and TCs conceptualised the experiments; TCs and AK designed the constructs and generated the transgenic lines; IA, RV and ISzK selected the plant lines and performed the RNA and protein works; HMSz and APSz performed protein alignment analysis; TCs wrote the draft with help of all authors in editing the final version of the manuscript; SD, ZH and TCs provided leadership and secured funding.

## Data availability Accession numbers

*HORVU*.*MOREX*.*r3*.*5HG0524690 (HvTFIIS*, Ensembl Plants, formerly *HORVU5Hr1G111700); A0A8I6YLM2 (HvTFIIS*, Uniprot*); Q9ZVH8 (AthTFIIS*, Uniprot*); AF-Q9ZVH8-F1 (AthTFIIS*, AlphaFold DB*); A0A317Y0Y7 (ZmTFIIS*, Uniprot*); AF-A0A317Y0Y7-F1 (ZmTFIIS*, AlphaFold DB*); A0A8J5L1I5 (ZoTFIIS*, Uniprot*); A0A3B6KPR6 (TaTFIIS*, Uniprot*); A3AP07 (OsTFIIS_A*, Uniprot*); Q0D7N2 (OsTFIIS_B*, Uniprot*); A0A9D5CLN9 (DzTFIIS*, Uniprot*); A0A0Q3N765 (BdTFIIS*, Uniprot*); A0A438J2H9 (VvTFIIS*, Uniprot*); A0A251SLR7 (HaTFIIS*, Uniprot*); A0A3Q7GBP4 (SlTFIIS*, Uniprot*); M1BUE0 (StTFIIS*, Uniprot*); A0A2G2ZJJ8 (CanTFIIS*, Uniprot*); A0A6P6W5C3 (CarTFIIS*, Uniprot*); A0A1S4C6W4 (NtTFIIS*, Uniprot*); A0A078F3V7 (BnTFIISa*, Uniprot*); A0A078FJ33 (BnTFIISb*, Uniprot*); A0A078JA64 (BnTFIISc*, Uniprot*); I1KB67 (GmTFIIS_1*, Uniprot*); C6TBG6 (GmTFIIS_2*, Uniprot*); I3SY13 (MtTFIIS*, Uniprot*); W1PB17 (AtTFIIS*, Uniprot*); XP_031484757 (NcTFIIS*, NCBI*); XP_058075383 (MsTFIIS*, NCBI*); A0A0D6R978 (AcTFIIS*, Uniprot*); A0A0C9S597 (WnTFIIS*, Uniprot*); A9NWY5 (PsTFIIS*, Uniprot).

