## Supplementary figures and tables supporting the work. for "TFIIS is required for reproductive development and thermal adaptation in barley": Fig-S1-S4-Table copy.pdf

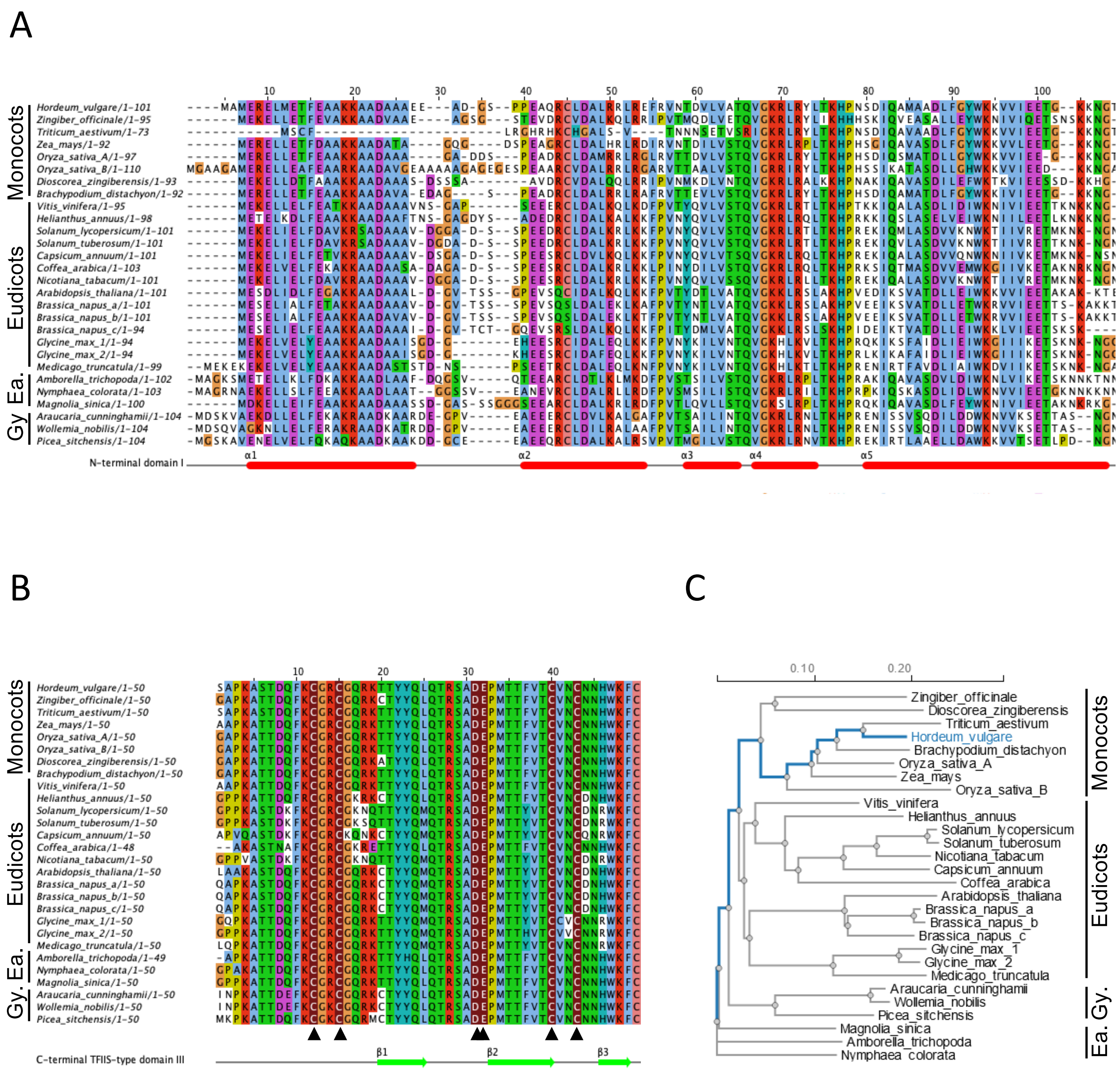

**Supplementary figure S1: Protein alignment of TFIIS protein homologs;** The sequences of gymnosperm (Gy), early dicot (Ea), eudicot and monocot TFIIS protein homologs (as shown on left) were aligned by Jalview ClustalO tool and secondary structure $\alpha$-helices and $\boldsymbol{\beta}$-sheets predicted (shown on bottom) for TFIIS domain I (A) or TFIIS domain III (B). The four cystein residues forming the zinc-finger and acidfic dipeptide of TFIIS domain III is shown with arrowheads below. (C) Phylogenetic tree of TFIIS homolgs in representative species of plant kingdom; the length of the branches is indicative for evolutionary distance between protein sequences.

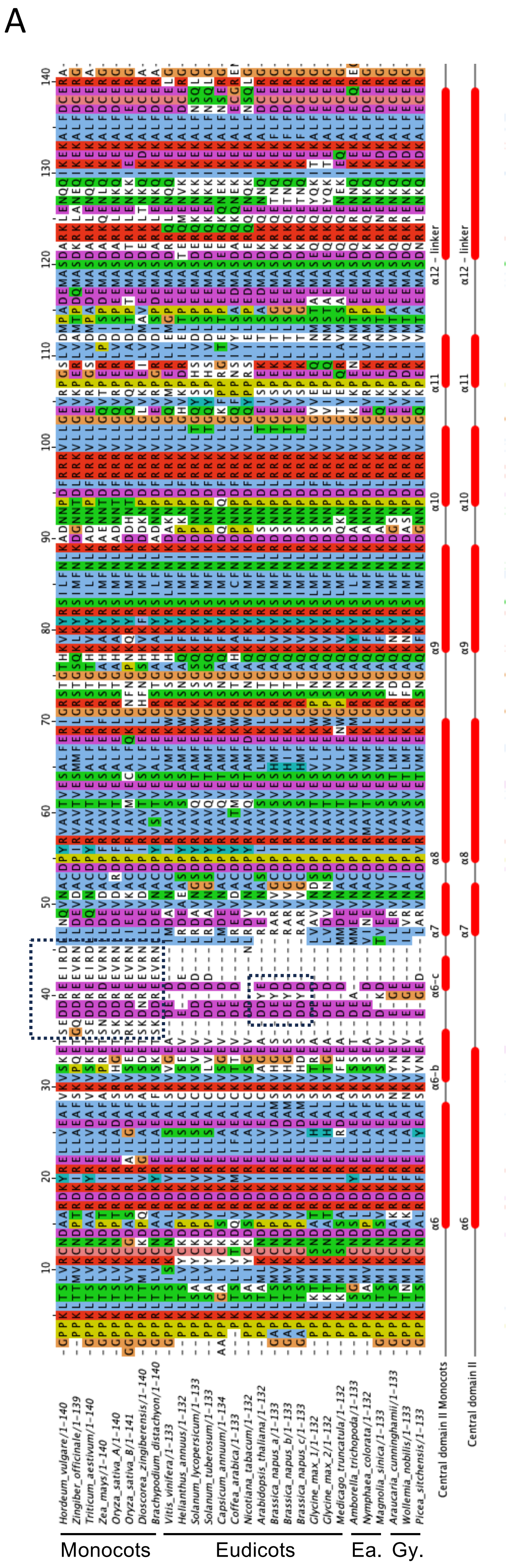

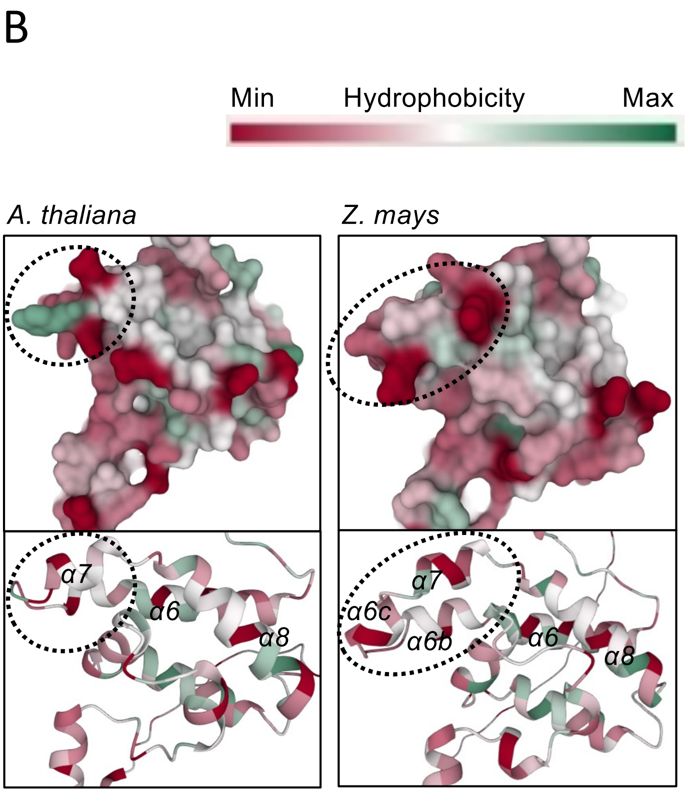

**Supplementary figure S2: Protein alignment of domain II of TFIIS protein homologs.** (A) The sequences of gymnosperm (Gy), early dicot (Ea), eudicot and monocot TFIIS protein homologs (as shown on bottom) were aligned by Jalview ClustalO tool and secondary structure $\alpha$-helices predicted (shown on the right) for TFIIS domain II. (B) The *in silico* 3D structure prediction of TFIIS domain II $\alpha$6-to-$\alpha$8 region of *A. thaliana* and *Zea mays* homologs; hydrophobicity heatmap color legend shown on top; the Arabidopsis-specific and Z. mays (monocot)-specific structural alterations are highlighted by dotted squares (A) and dotted circles (B).

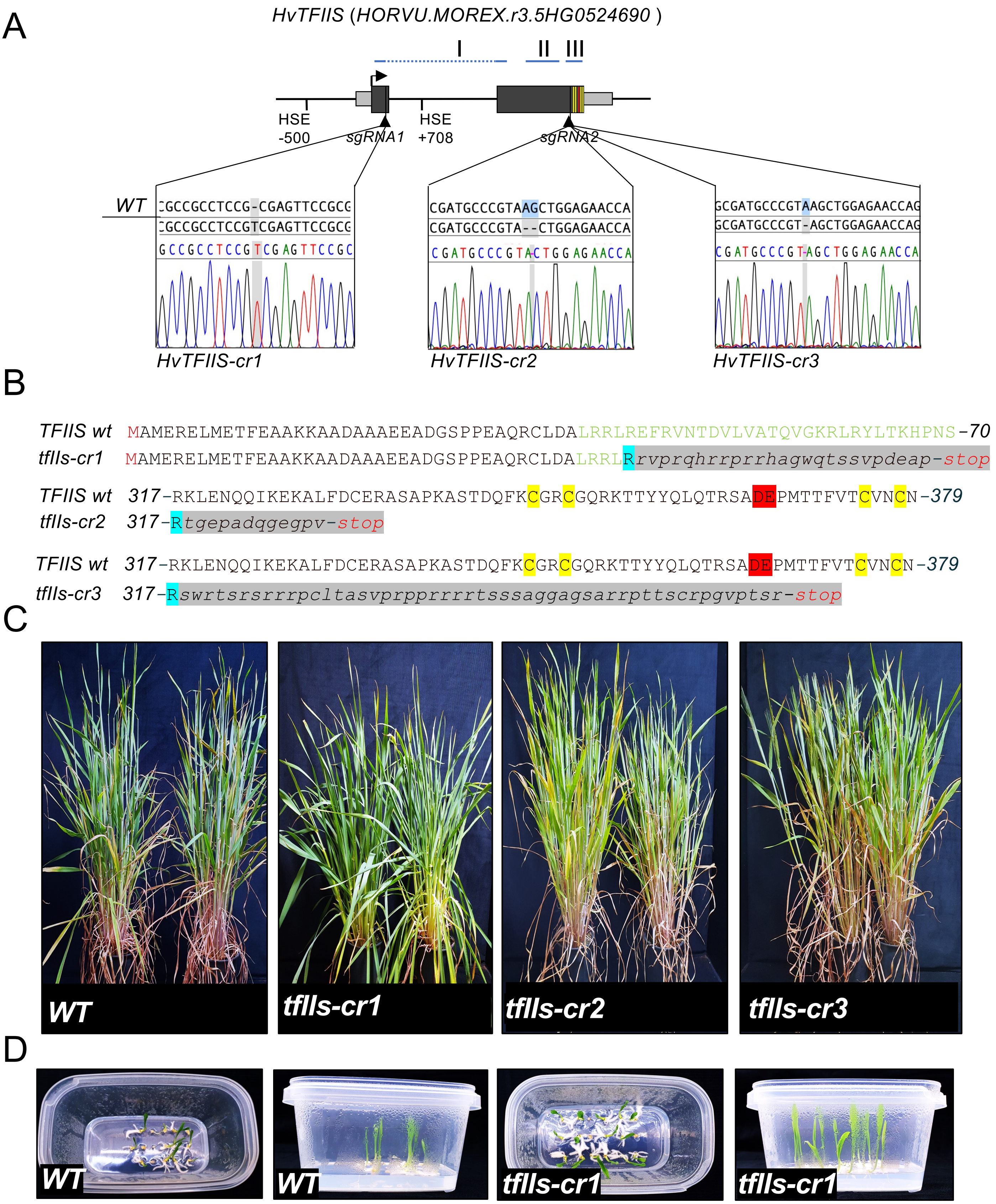

**Supplementary figure S3: TFIIS is negligible for vegetative growth but needed for reproductive development in barley.** (A) (A) Schematic representation of *HvTFIIS* gene locus; protein domain I, II and III location is shown above; exons as black boxes, UTRs as grey boxes, Zn finger domain cysteine residues and DE acidic dipeptide motif as yellow and red line, respectively; HSE *cis* elements and location of sgRNA guide sites are shown below; (B) TFIIS mutants of barley were created by CRISPR mutagenesis; Insertion or deletion mutation within the TFIIS locus in the selected transgenic lines are shown on chromatograms; (C) HvTFIIS protein amino-acid changes with premature termination codon is shown below; last correct aminoacid is highlighted with cyan, changed amino acid sequence with grey, Zn-finger motif residues with yellow, catalytic DE dipeptide with red; Vegetative growth of wild tpye (wt) and CRISPR mutant *tfIIs-cr1*, *-cr2* and *-cr3* barley plants; *tfIIs-cr1* plants are slightly shorter in stature, while *tfIIs-cr2* and *-cr3* plants are indistinguishable to wt plants; (D) Embryo rescue of *tfIIs-cr1* seeds, wild-type and mutant embryos were extracted from the seeds and placed on sterile germinating agar medium for growth; wt and mutant embryos were treated similarly for generation of plants and TMHT treatment work shown in Fig 2 and S4.

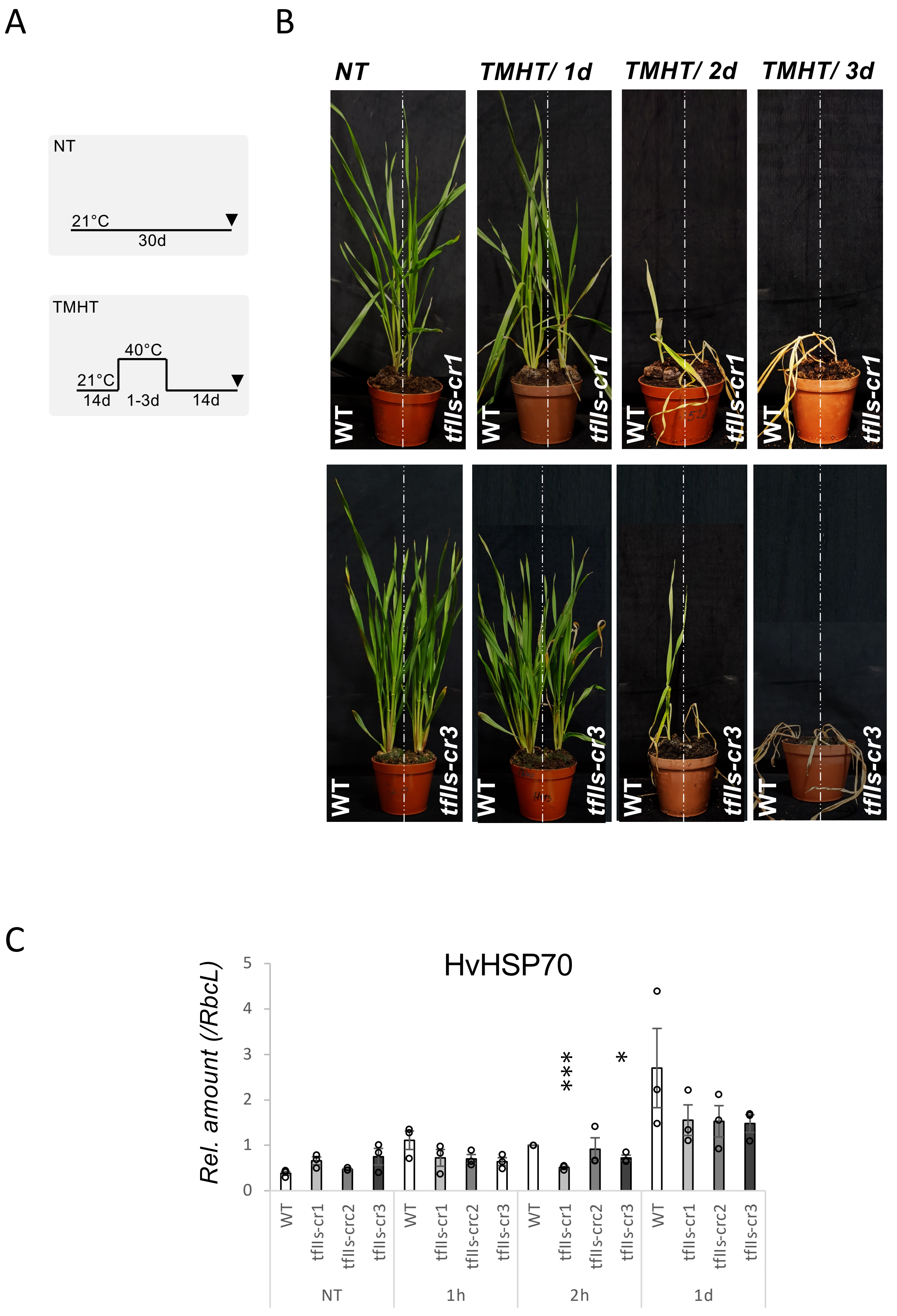

**Supplementary figure S4: TFIIS is indispensable for efficient heat stress response.** (A) Schematic representation of non-treated (NT) and Thermotolerance to Moderately High Temperature regime (TMHT); arrowheads show the time of sampling; (B) Wild-type (wt) and *tfIIs* CRISPR mutant *tfIIs-cr1* (top) and *tfIIs-cr3* (bottom) were exposed to NT or TMHT for 1 hour (1h), 2h or 1 day (1d); (C) Quantification of HSP70 protein amount changes in response to heat stress during a timeseries; bars represent standard errors based on three bio reps; P-values based on two-tailed Student's t-test (*P < 0.05, **P < 0.01, ***P < 0.001) show differences between wild type and mutant plants.

**Supplementary Table 1: DNA oligonucleotide primers used in the study.**

| | |
|---|---|
| Crispr constructs | |
| HvTFIIS_cr1_F | GGC GGT TGA CGC GGA ACT CGC GG |
| HvTFIIS_cr1_R | AAA CCC GCG AGT TCC GCG TCA AC |
| HvTFIIS_cr2_F | GGC GGC GAG CGA TGC CCG TAA GC |
| HvTFIIS_cr2_R | AAA CGC TTA CGG GCA TCG CTC GC |
| Genotyping | |
| HvTFIIS_crispr1_check_F | CTG CGC CTA GGG TTT TCA TC |
| HvTFIIS_crispr1_check_R | GCT TTC CGC ACC TCT CAG TC |
| HvTFIIS_crispr2_check_F | ACA GTG GAG TCG GCC TTG TT |
| HvTFIIS_crispr2_check_R | TAC GAA AAA CCG AGC CAA CC |
| HvTFIIS_cr1_check-gF | GGA GAG GGA GCT GAT GGA GA |
| HvTFIIS_cr1_check-gR | CGA TAG GGC ACG GAT AGT CTG |
| qrtPCR | |
| HvTFIIS_qF: | TCG CCA CGC AGG TTG GCA AAC G |
| HvTFIIS_qR: | TTC AAT AAC AAC CTT CTT CCA G |
| HvActin7_qF | CGT GTT GGA TTC TGG TGA TG |
| HvActin7_qR | AGC CAC ATA TGC GAG CTT CT |
| HvHSPc70-4_qF | CAA CAC CGT TTT TGA TGC CA |
| HvHSPc70-4_qR | ACC ACG ATC ATC GGC TTG T |
| HvHSP90_qF | AAC TCA TCT GAC GCG CTT GA |
| HvHSP90_qR | GCT GTC GAT GAT GGA GAG CGT |
| HvHSP101-1_qF | GTC ATG CAG GAG GTG AGG AGG |
| HvHSP101-1_qR | CCA CGT CCT TCA TCT GCA G |
